# Fall risk-aware adaptation explains suboptimal locomotor performance

**DOI:** 10.64898/2026.03.02.709033

**Authors:** Inseung Kang, Kanishka Mitra, Nidhi Seethapathi

**Affiliations:** Department of Brain and Cognitive Sciences, Massachusetts Institute of Technology, Cambridge, MA, USA; Department of Electrical Engineering and Computer Science, Massachusetts Institute of Technology, Cambridge, MA, USA; Department of Mechanical Engineering, Carnegie Mellon University, Pittsburgh, PA, USA

**Keywords:** Locomotion, Motor Performance, Energetics, Stability, Motor Learning

## Abstract

Human locomotion requires balancing multiple biological objectives, such as metabolic energy efficiency, stability, and symmetry. While models based on optimization successfully predict how humans walk in familiar settings, they fail to explain why individuals adopt inefficient movement patterns in novel environments, even after extensive practice. Here, we show that such suboptimality in a novel environment arises from a fundamental prioritization of safety. We find that individuals do not simply fail to reach an optimal solution; instead, they navigate an environment-dependent risk landscape by mitigating the statistical probability of falling. We find that this risk-averse strategy is explained by adjusting internal learning parameters: specifically, the learning rate and the tradeoff between metabolic cost and symmetry, in a manner that lowers fall risk. To quantify this process, we developed an ‘inverse adaptation’ modeling framework; this approach works backwards from locomotor performance data to mathematically infer the underlying internal learning parameters and how they vary with fall risk. Our analysis reveals that the observed motor performance is explained by a global probabilistic fall risk rather than a local step-based measure of instability. Ultimately, these findings reveal that fall risk-aware adaptation explains suboptimal locomotor behavior, providing a new data-driven framework to understand the drivers of motor performance.

## 1 Introduction

In everyday life, humans continually adjust their locomotion while negotiating multiple, often competing, objectives including energy efficiency [1, 2], safety and stability[3, 4], and symmetry [5, 6]. Despite this apparent flexibility, the principles that govern how individuals navigate multiple goals, particularly when exposed to unfamiliar environments, remain poorly understood. In addition to posing a fundamental scientific gap, this limitation constrains the development and effective adoption of assistive technologies such as rehabilitation robots and wearable exoskeletons [7, 8], which frequently constitute novel environments for users. Although computational models grounded in optimization frameworks have advanced our understanding of how performance goals shape motor behavior [9, 10], they typically assume rapid and idealized convergence toward an optimal solution even in a novel environment [11, 12], not accounting for the gradual or energetically suboptimal performance observed in practice [13, 14]. These assumptions restrict our ability to explain how motor performance unfolds over time and limit the interpretive and practical relevance of existing modeling approaches [15, 16]. To address this, we introduce an ‘inverse adaptation’ modeling paradigm that leverages time-varying behavioral data in a novel environment to understand the drivers of suboptimal performance.

A central challenge in interpreting motor performance lies in understanding the interplay between multiple objectives like energy efficiency, symmetry, and stability. Although these objectives co-exist in reality, most studies examine them in isolation, which can obscure how performance emerges from their interaction. This isolation results in apparent suboptimality along an individual objective; for example, the failure to reach an energetic optimum in some settings [17, 18] and the varying degrees of symmetry observed in others [19, 13] are well-documented, yet the principles governing these deviations remain unclear. One possible explanation for such apparent suboptimality is that the nervous system may prioritize other performance goals—such as maintaining stability or safety—in response to environmental demands. As multiple performance goals can interact, experimental approaches alone struggle to capture their combined influence due to the inherent combinatorial complexity of the problem. Computational modeling in combination with experimental data therefore provides a necessary framework for decoupling these interactions and identifying the principles that govern adaptive motor performance.

Computational models based on optimization have yielded important insights into human locomotion, accounting for features as preferred speed [2, 20], step frequency [21], and navigation trajectories [22]. However, their ability to explain behavior in unfamiliar environments remains limited; while these models can identify energetic optima in novel conditions [23, 24, 12], an assumption of consistent convergence prevents them from explaining important observations, such as why performance remains suboptimal relative to predictions [17, 18] or why performance converges at different rates [13, 14]. Critically, classic trajectory optimization relies on periodicity constraints that preclude falls in simulation [25, 26]. Lacking the ability to simulate fall events or feedback control deficits, these models cannot effectively test interactions between stability and other goals. While recent work has modeled adaptation as an interaction between energy and symmetry goals within a framework capable of falling [27], we lack a framework for understanding how these goals are reweighed in response to environmental demands. Here, we put forth an ‘inverse adaptation’ modeling framework to infer the learning parameters underlying individual performance, finding that the mitigation of fall risk explains seemingly suboptimal energetic performance.

In this work, we discover that suboptimal locomotor performance in a novel environment is explained by fall risk-aware adaptation. To do this, we put forth an ‘inverse adaptation’ modeling paradigm that infers how learning parameters that impact locomotor performance are modulated in response to environmental fall risk. We choose split-belt walking as a novel environment and model system, due to its documented effects on multiple motor objectives, including energy efficiency, symmetry, and stability [13, 28]. By applying our framework to time-varying individual locomotor data, we estimate how learning rates and objective weights shift in response to the environmental setting—namely the belt speed difference—explaining environment-dependent performance shifts. Crucially, we find that these seemingly suboptimal performance shifts are explained by a strategy that prioritizes safety through the mitigation of fall risk. Overall, this work offers a computational basis for evaluating how safety constraints and motor objectives interact to shape environment-dependent motor performance.

## 2 Results

We examine human locomotor behavior across multiple performance objectives—energy, symmetry, and fall risk—within a novel environment. Because split-belt walking simultaneously influences these interacting objectives [29, 30, 31], it serves as a suitable experimental platform for probing how locomotor performance unfolds. We study how performance varies with exposure to the split-belt environment across different settings, namely belt speed differences. We find that our ‘inverse adaptation’ framework more accurately captures observed locomotor performance across settings compared to optimization-based model predictions. By aligning time-varying performance with simulated adaptation profiles, this framework identifies the internal learning parameters—specifically the learning rate and the energy-symmetry tradeoff—that best characterize individual performance. Furthermore, by quantifying probabilistic fall risk across a range of learning parameters for each setting, we find that this risk explains the parameters selected by individuals, revealing a fundamental relationship between environment-dependent fall risk and adaptive performance.

### 2.1 Time-varying locomotor performance in a novel environment is not explained by energy minimization

#### Split-belt walking as a model system for examining environment-dependent locomotor performance

Under-standing how locomotor performance evolves in a novel environment and varies with environmental settings is a central scientific question with direct implications for the design of rehabilitation technologies. Modifying the belt speed difference or the amount of exoskeleton assistance fundamentally alters the energetic, symmetric, and stability performance experienced by the user [13, 14, 33, 34, 31, 35]. In this work, we use split-belt walking as a model system to systematically probe how changes in the environmental setting—namely the belt speed difference—influence multiple dimensions of locomotor performance. This paradigm is particularly well suited for this question because it perturbs key locomotor objectives concurrently; much like exoskeletons, split-belt treadmills can deliver positive mechanical work to the user, reducing metabolic energy expenditure [36, 37]. While their influence on gait symmetry is well established [19, 30], evidence also suggests a consequential impact on stability [31, 27]. Together, these properties position split-belt walking as an ideal experimental platform for understanding the interaction between environmental demands and locomotor performance.

#### Energy minimization does not explain environment-dependent locomotor performance

To evaluate how locomotor performance unfolds across different settings in a novel environment, we systematically varied the belt speed difference on a split-belt treadmill (Figure 1A, B). Metabolic energy expenditure and step length asymmetry were quantified over a 45-minute period for each condition; percentage changes were assessed relative to tied-belt walking across both short (6-minute) and long (45-minute) timescales. To verify the reliability of our measurements, we compared our energetic and symmetry findings in the high belt speed setting to those of [32], the only setting examined in that study. Our measurements were quantitatively consistent with their findings for both 6 and 45 minutes of exposure, confirming that these observations are replicable.

**Figure 1.**
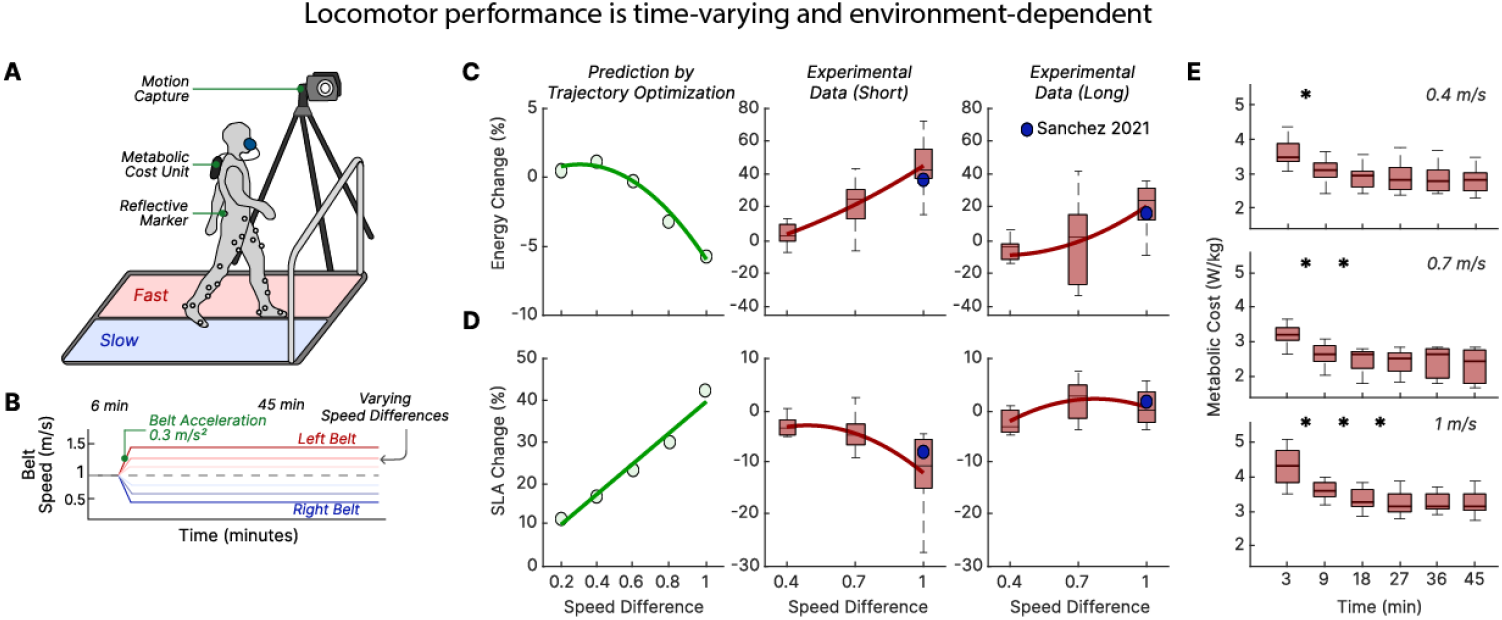
Locomotor performance is time-varying and environment-dependent. **A**. Experimental setup: Participants walked on a split-belt treadmill, where both belt speeds can be independently controlled, while whole-body kinematics and indirect calorimetry were recorded. **B**. Belt speed differences defined three environmental settings: low (0.4 m/s), moderate (0.7 m/s), and high (1.0 m/s). Comparison of optimization-based model predictions and experimentally observed **C**. metabolic energy change and **D**. step length asymmetry. The trajectory optimization predictions (leftmost, green generated using [12]) are compared against observed behavior (right, pink) over short (6 min) and long (45 min) exposure. This comparison highlights discrepancies between energetic optimality-based model predictions and the observed performance across timescales. Long-duration performance data on a split-belt treadmill from prior work in a single setting [32] are included (blue). **E**. Temporal evolution of metabolic cost across environments, illustrating that the timescale of energy minimization depends on the environmental setting. Box plots show inter-subject distributions; asterisks indicate significant differences between consecutive time intervals.

These empirical observations were then compared against predictions from an energy minimization model generated by a trajectory optimization-based musculoskeletal framework [12]. This model predicts that energy expenditure should decrease monotonically with increasing belt speed difference, implying that larger speed disparities provide a mechanical advantage consistent with prior theoretical work [23, 38]. In contrast, experimental measurements revealed qualitatively divergent behavior across both timescales compared to model predictions; rather than decreasing, metabolic cost increased with greater belt speed differences (Figure 1C). Step length asymmetry similarly deviated: while the model suggested increasing asymmetry, participants exhibited decreasing asymmetry over short timescales and near-zero asymmetry over prolonged exposure (Figure 1D). These discrepancies indicate that energy minimization alone does not account for observed locomotor performance, suggesting that alternative performance goals drive motor behavior in novel environments.

#### The timescale of energy minimization is environment-dependent

The timescale over which motor objectives like metabolic energy converge is an important feature of locomotor performance in a novel environment. Although many assistive strategies might aim to reduce metabolic energy quickly upon exposure to novel settings, such reduction unfolds over extended time periods in practice [14, 13]. While such modulation of the learning rate has been well characterized in error-based motor learning [39, 40], the environment-dependent timescale over which other motor objectives, like energy minimization, unfold remain underexplored. Prior work has demonstrated that prolonged exposure to split-belt walking can yield reductions in energy expenditure [29, 32], yet how the timescale of such reduction varies with belt speed difference has not been studied. Metabolic cost in this study exhibited a rapid initial decrease across all conditions during the first 10 minutes (Figure 1E), followed by different timescales of decrease across environmental settings. Under low belt speed differences, metabolic cost converged quickly, showing minimal further change. In contrast, higher speed differences produced a slower decline in metabolic cost, extending through approximately half of the session. These findings demonstrate that the timescale of energy minimization is dependent on the environmental setting. These distinct timescales of energy minimization further suggest that alternative performance objectives might underlie environment-dependent motor performance.

### 2.2 Inverse adaptation model captures time-varying locomotor performance across environmental settings

We developed an inverse adaptation model by leveraging a recent forward model of locomotor adaptation [27] that integrates a low-level feedback controller with a high-level policy-gradient reinforcement learner (Figure 2). Two key parameters govern the learner’s trajectory: the learning rate *α*, determining the rate of policy updates, and the symmetry weight *ψ*, determining the prioritization of symmetry relative to energy expenditure. This approach identifies the internal parameters of the learner by mapping experimental energy trajectories to a pre-computed library of forward simulations (Figure 3). For each individual, we resolve the specific combination of *α* and *ψ* that minimizes the discrepancy between the simulated and observed performance. This approach allows us to recover the internal learning parameters that characterize individual-specific responses to each environmental setting. The resulting fits reveal close correspondence between model predictions and observed performance, with RMSE values of 0.33 ± 0.32%, 0.90 ± 0.83%, and 1.28 ± 0.49% for the 0.4, 0.7, and 1.0 m/s belt speed difference settings, respectively. Relative to the magnitude of total energy change over a trial, these low error values indicate that the inverse adaptation model robustly captures the time-varying structure of individual locomotor performance across settings.

**Figure 2.**
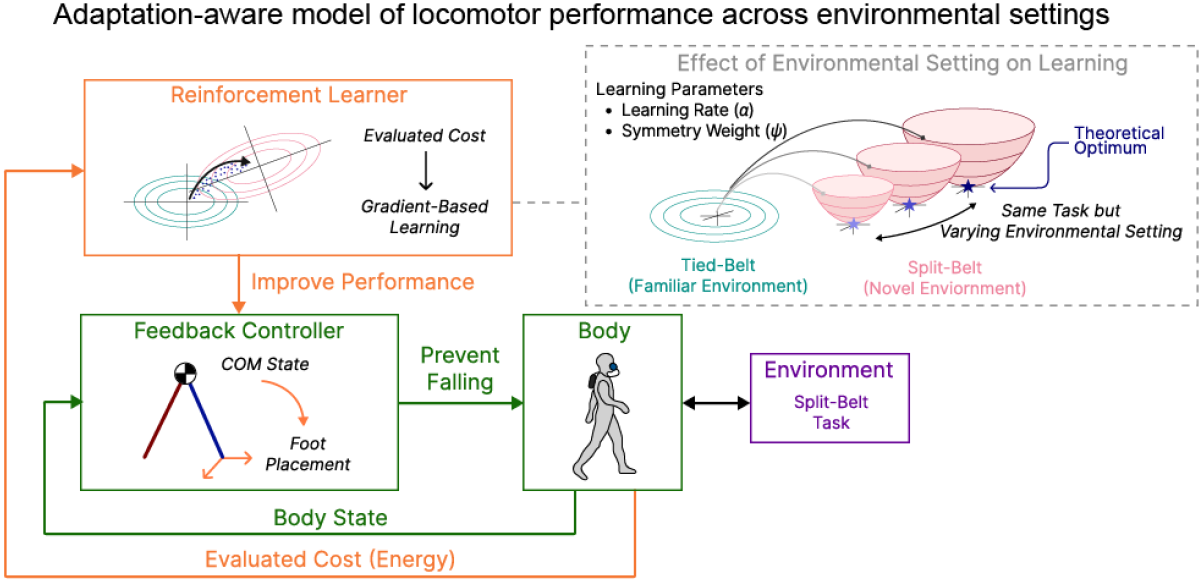
Adaptation-aware model of locomotor performance across environmental settings. The model, adapted from [27], comprises a hierarchical structure in which a low-level feedback controller (green) determines step-by-step foot placement based on body-state feedback to maintain stability. In novel locomotor environments (purple), such as split-belt walking, a high-level reinforcement learner (orange) modifies the control policy using gradient-based updates to improve performance. Two key parameters govern this adaptation process: the learning rate, which modulates the rate of policy updates, and the symmetry weight, which modulates the prioritization of symmetry relative to energy minimization. The environmental setting (dashed box) is varied, inducing distinct theoretical optima and time-varying adaptive performance.

**Figure 3.**
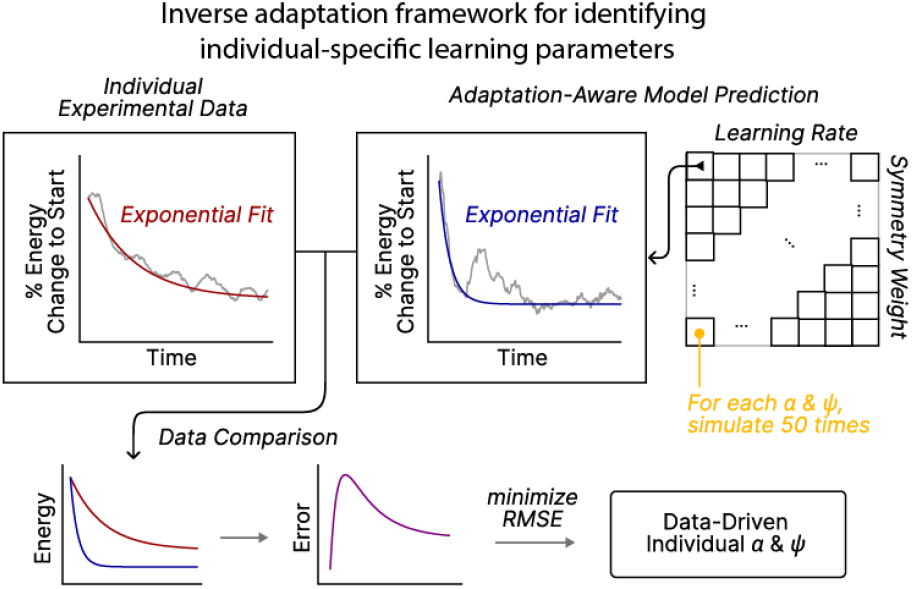
Inverse adaptation framework for inferring individual-specific learning parameters. The inverse adaptation model integrates experimental data with model-based simulations of locomotor adaptation (Figure 2) to identify the learning parameters governing individual locomotor performance. For each trial, a first-order exponential function was fitted to the experimentally measured time-varying energy profile. In parallel, the model was simulated across a range of learning rates and symmetry weights to generate a library of simulated energy profiles, also each fitted with a first order exponential. The parameter combination that minimized the root-mean-square error (RMSE) between the experimental and simulated profiles was selected as the best-fit representation of each individual’s adaptive performance.

### 2.3. The probabilistic fall risk landscape explains environment-dependent locomotor performance

#### Fall probability quantifies how adaptation influences instability

A defining feature of the physics-based simulation (Figure 4A) we use in our inverse adaptation model is its capacity to simulate falls, in contrast to trajectory optimization approaches that impose periodicity constraints, thereby precluding instability. Because the model incorporates step-by-step feedback control, certain parameter choices can result in deviations in the center-of-mass trajectory, resulting in falls. These events were identified when a sudden vertical drop occurred during a simulated step (Figure 4B). To understand how the model influences stability, we quantified two measures: (i) probabilistic fall risk and (ii) the number of strides before a fall, analyzing how these measures vary with the selected learning rate *α*. We found that, across all speed difference settings, fall probability increased systematically with the learning rate (Figure 4C), and escalated further as the belt speed difference increased. This indicates that the probabilistic risk of falling is sensitive to both the learning rate and the environmental setting. In contrast, the number of strides before a fall exhibited substantial variability and a much weaker relationship with the learning rate (Figure 4D), indicating that probabilistic fall risk is more influenced by the selection of learning parameters and is a more sensitive measure of environment-dependent instability. Taken together, these results reveal an intrinsic tradeoff: higher learning rates benefit energy minimization but simultaneously amplify fall risk under certain environmental settings.

**Figure 4.**
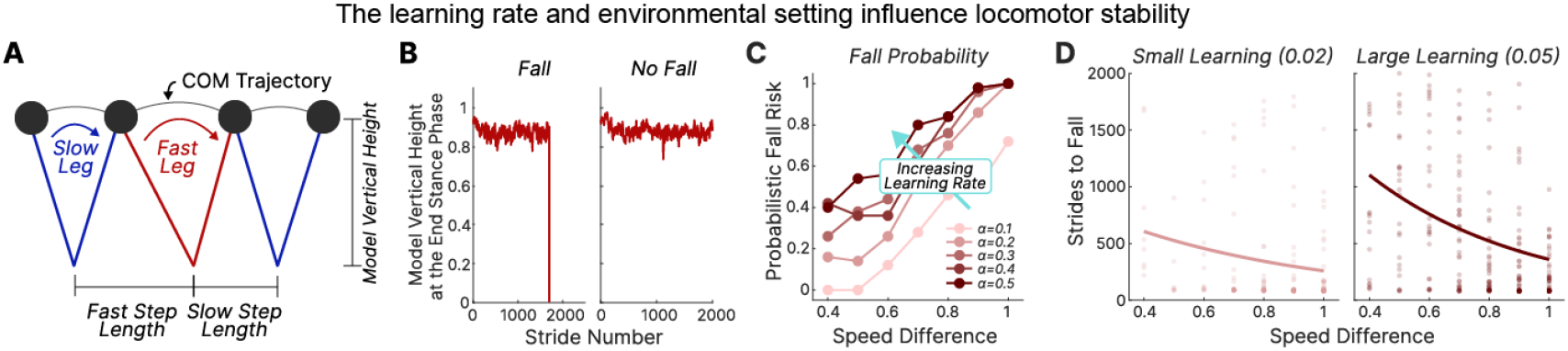
The learning rate and environmental setting influence locomotor stability. **A**. Representative locomotor pattern exhibited by the model during stable gait. **B**. Example simulations illustrating instability during adaptation. Vertical center of mass (COM) height at the end of stance is shown for a simulation that results in a fall (left) compared to one that remains stable (right). **C**. Simulated fall probability as a function of learning rate *α* across environmental settings, demonstrating a systematic increase in fall risk with both higher learning rates and larger belt speed differences. **D**. Number of strides until a fall event occurs across learning rates and environmental settings, overlaid with best-fit exponential curves. Each dot represents an independent simulation.

#### Learning parameters are modulated across environmental settings in a fall risk-averse manner

Inverse adaptation revealed that best-fit learning rates declined with increasing belt speed differences, while symmetry weights increased correspondingly. We find that this pattern is well explained by the environment-dependent fall risk landscape (Figure 5). Indeed, maintaining the same parameter combinations across different environmental settings would result in significantly increased fall probability (*p <* 0.05), as illustrated by the divergence between the grey and red ellipses in Figure 5. This highlights the necessity of environment-dependent modulation.

**Figure 5.**
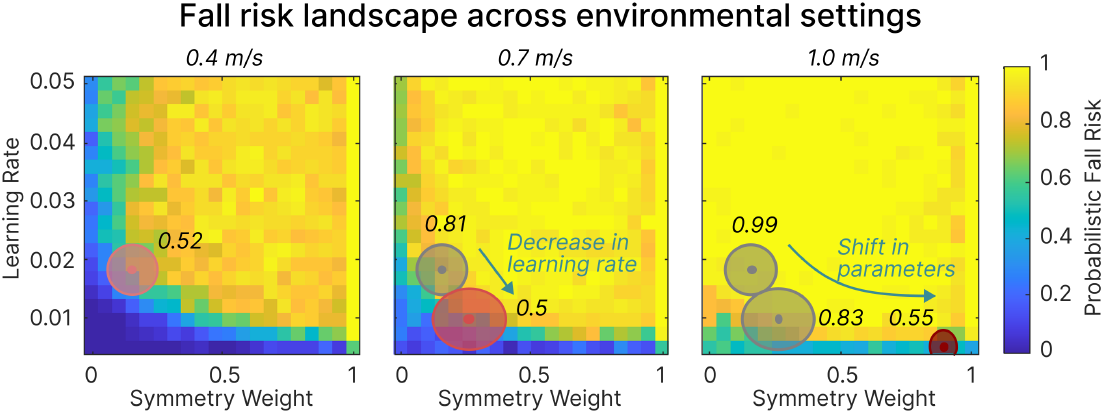
Fall risk explains the modulation of learning parameters across environmental settings. Simulated fall risk landscapes are shown across environmental settings as a function of learning rate and symmetry weight, the key parameters influencing locomotor performance. Each heatmap represents the probabilistic fall risk across the parameter space for a specific belt speed difference. Ellipses denote population-level parameter regions inferred through inverse adaptation; red ellipses correspond to the specified setting, while grey ellipses represent settings with lower belt speed differences. Centers of the ellipses represent the median learning rate and symmetry weight across individuals, with major and minor axes denoting ± standard error. Although the 1.0 m/s ellipse appears to extend below the axis, all simulated learning rates were constrained to values greater than 0.005. Percentages adjacent to each ellipse indicate the mean fall probability within the encompassed parameter space. As indicated by the overlaid arrows, individuals shift toward lower learning rates and higher symmetry weights as the landscape risk increases, strategically occupying safer regions of the parameter space.

Overall, mapping subject-specific parameter distributions onto the fall risk landscape revealed a consistent population-level shift toward regions characterized by lower learning rates and higher symmetry weighting as environmental challenge increased; these shifts are consistent with occupying regions of lower fall probability (Figure 5). Furthermore, the variability in parameter distributions across individuals narrowed at the highest speed difference, mirroring the shrinking regions of safe operation at this setting. This indicates that individuals converge toward a constrained parameter range in environments with high fall risk. Collectively, these observations indicate that locomotor adaptation is governed not primarily by improvements in energy or symmetry, but by the avoidance of fall risk. Specifically, individuals modulate their learning rate and symmetry weight across environments in a fall risk-averse manner.

## 3 Discussion

In this work, we introduced a data-driven, model-guided framework for understanding how learning parameters are selected across settings, thereby influencing locomotor performance. Using this framework, we discovered that individuals select their learning rate and symmetry weight in novel walking environments in a manner that reduces fall risk. This ‘inverse adaptation’ framework identifies individual-specific learning parameters by aligning time-varying model performance with each individual’s experimental behavior. A key contribution is the use of simulated fall probability to quantify fall risk across candidate learning parameters, explaining why individuals exhibit performance trends that cannot be explained by energy minimization or symmetry alone. Collectively, these findings advance our understanding of the interplay between energy efficiency, stability, and symmetry, providing quantitative insight into how these objectives interact to shape motor performance over time.

### Inverse adaptation reveals the drivers of motor performance in a novel environment

Understanding the factors that drive motor performance in novel environmental settings remains a fundamental challenge that current models do not satisfactorily address. In this study, we propose an inverse adaptation modeling paradigm that combines forward simulations with data-driven inference to identify the learning parameters that best explain each individual’s time-varying performance. A prominent alternative, trajectory optimization, posits that humans select movements that optimize a specific objective—such as minimizing metabolic energy or asymmetry—allowing for comparisons between theoretically optimal solutions and human behavior [24, 12, 23, 25]. However, trajectory optimization assumes complete, instantaneous optimization and fails to account for the time-varying [13, 14] or seemingly suboptimal performance typically observed in novel environments [17, 18]. Indeed, we demonstrate that energy minimization-based trajectory optimization fails to capture observed variations in locomotor performance across environmental conditions, even at a qualitative level (Figure 1). Inverse reinforcement learning (IRL) is another data-driven method used to explain the objectives underlying observed behavior by quantifying a reward function from policy rollouts [41, 42, 43]. Yet, standard IRL models presume expertise by assuming that the observed behavior is already optimized for a cost function; this limits their applicability in novel or destabilizing environments—frequently encountered in rehabilitation and wearable robotics—where assumptions of optimality may not hold. In contrast, our inverse adaptation framework avoids the assumption of perfect convergence toward a theoretical optimum. Instead, it explicitly models the adaptation process to recover the individual-specific learning parameters underlying the observed time-varying performance.

### In silico fall probability landscapes to quantify environment-dependent risk

Our environment profoundly influences how we move, with perceived instability playing a critical role in shaping motor behavior [44, 45]. However, existing models offer limited insight into how environment-dependent fall risk guides movement strategies, particularly in terms of adaptive performance in novel settings. To address this, we introduce the concept of a “fall risk landscape” as a parallel to the classic “energy landscapes” used to map the metabolic cost of movement [1, 21, 22]. In our framework, this landscape explains how environment-dependent fall risk drives individuals to exhibit adaptive performance (Figure 5). Further, we found that the number of strides before a fall—a common metric for assessing instability in silico [46, 47, 48]—was not a reliable indicator of environment-dependent risk due to its high variance and weak sensitivity to learning parameters (Figure 4D). In contrast, quantifying probabilistic fall risk through simulation revealed a clear relationship between learning parameters and instability; we found that individuals adopt learning rates and symmetry weights that maintain their adaptive performance within low-risk regions of this landscape. By defining stability through the lens of fall probability, our work advances a perspective on motor control that emphasizes the proactive avoidance of instability [49]. This suggests that the nervous system does not simply react to instability as it occurs, but instead proactively modulates adaptive performance to avoid high-risk parameter combinations, highlighting the importance of probabilistic, environment-sensitive measures in addition to widely used step-based metrics.

### New insight into how fall risk influences the timescale of energy minimization

We found that individuals modulate their learning rate and symmetry weighting across environments in a manner that mitigates fall risk. Specifically, in settings with higher fall risk participants exhibited lower learning rates, slower reductions in metabolic cost, and higher weighting of symmetry relative to energy. This pattern indicates a shift toward more conservative motor strategies in the presence of higher fall risk. These findings may reconcile the variable energy minimization timescales reported previously [50, 51, 17, 32, 14], suggesting that prolonged timescales of adaptive performance may reflect environments associated with increased fall risk. Our simulations further reveal that aggressively pursuing energy minimization in high-risk conditions can destabilize gait. In such scenarios, individuals may intentionally limit the rate of adaptive performance or increase symmetry weighting to prioritize stability, demonstrating that fall risk can directly influence the pursuit of energetic efficiency in a novel environment. These findings indicate that preserving balance can supersede energy minimization in some environments, thereby extending the timescale of performance improvement.

### Gait symmetry as a means to reduce the risk of falling

Our findings provide new insight into the role of gait symmetry during locomotor learning as a strategy for mitigating fall risk. While previous work suggested that individuals might retain asymmetric patterns over long timescales to reduce metabolic cost [13], the asymmetries we observed here (Figure 1D) were substantially lower than those predicted by energy minimization alone [23, 12, 27]. Indeed, in contrast to predictions derived solely from energy minimization, we observed a trend toward symmetry over extended timescales, particularly under larger belt speed differences. In agreement with these experimental findings, our inverse adaptation model reveals that individuals increase their symmetry weighting in higher-risk environments, thereby avoiding fall risk (Figure 5). We note that the literature on locomotor asymmetry remains mixed, with some studies reporting a preference for symmetry [33, 52] and others identifying persistent asymmetry [13, 32]. Our results suggest that these discrepancies may arise from variations in real or perceived fall risk across experimental settings. From this perspective, the degree of observed asymmetry may reflect a fall risk-averse strategy rather than an inherent preference for symmetry itself.

### Limitations and future work

Our framework models human locomotion using low-dimensional physics governed by a structured feedback controller. While this approach efficiently captures the fundamental aspects of adaptive behavior, it serves as a baseline that does not explicitly account for more complex recovery mechanisms, such as stumbling-recovery strategies [53] or upper-body compensations [54]. Future research should incorporate detailed neuromechanical models and recovery dynamics to further refine estimates of environment-dependent fall probability. In this study, we utilized a split-belt locomotion task because it probes multiple motor objectives simultaneously and provides a reproducible baseline using widely available hardware. Future work should extend this approach to other environmental contexts, such as assistive exoskeletons and prostheses [14, 18], enabling the fall risk-aware personalization of device control strategies. Additionally, exploring how model-inferred trade-offs correspond to subjective perceptions of stability and comfort could clarify how individuals internally prioritize safety relative to energetic efficiency [55].

### Conclusions

This work introduces an inverse adaptation modeling paradigm that explains the drivers of motor performance across environmental settings and accounts for time-varying, seemingly suboptimal behavior. Our results demonstrate that motor performance reflects the interplay between multiple objectives—including energetics, symmetry, and a probabilistic measure of fall risk—rather than a single optimization target. We show that environment-dependent fall risk exerts a fundamental influence on both the timescale of metabolic energy minimization and the degree of gait symmetry. These findings underscore the necessity of using adaptation-aware modeling paradigms, rather than assuming instantaneous optimality, to characterize human motor performance. Further refinement and extension of this inverse adaptation approach hold promise for advancing our understanding of adaptive performance and for informing the design of personalized, safety-aware assistive and rehabilitation technologies.

## 4 Materials and Methods

We combined human subject experiments (Figure 1A,B), model-based simulations (Figure 1C,D; Figure 2), and a data-driven, model-guided framework that we term “inverse adaptation” (Figure 3) to investigate locomotor performance across multiple locomotor objectives in a novel environmental setting. We conducted split-belt treadmill experiments in which the two belts moved at different speeds and quantified how subjects’ energy expenditure and gait symmetry evolved as they adapted to these speed-difference settings. To interpret these experimental findings, we used two complementary computational approaches: (1) an adaptation-aware model embedded within an inverse adaptation framework, and (2) a benchmark trajectory optimization model. In the inverse adaptation setup, we inferred individual- and environment-specific learning parameters by matching the time-varying experimental performance to simulated adaptation trajectories. Using the same model, we then quantified where the inferred learning parameters lie on a fall risk landscape derived from simulated fall probabilities. Both Python and MATLAB implementations of the inverse adaptation model (repository name: *InvLocAd*) will be shared openly upon publication.

### 4.1 Measuring locomotor performance across environmental settings and timescales

#### Experimental protocol

We recruited 27 healthy young adults (7 females; age: 22.5 ± 3.4 years; height: 1.73 ± 0.11 m; body mass: 74.1 ± 17.5 kg). None of the participants had prior experience walking on a split-belt treadmill, minimizing potential confounds from pre-existing adaptation. All experiments were performed on an instrumented split-belt treadmill (Bertec, Ohio). A safety handrail was available for all trials; participants were instructed to use it only if absolutely necessary to reduce its influence on natural gait. The study protocol was approved by the Institutional Review Board with protocol number 2012H0032, and written informed consent was obtained from all participants prior to testing. Each participant completed a single experimental session consisting of two continuous walking conditions (Figure 1B): (1) a 6-minute tied-belt condition at 1.0 m/s, followed by (2) a 45-minute split-belt condition. In the split-belt condition, participants were randomly assigned to one of three belt speed difference settings: 0.4 m/s, 0.7 m/s, or 1.0 m/s, while maintaining an average belt speed of 1.0 m/s across all groups. For the split-belt phase, the left belt always operated at the higher speed and the right belt at the lower speed, ensuring a consistent mapping between fast and slow sides across subjects.

#### Data acquisition

Metabolic energy expenditure was measured using indirect calorimetry (K5, COSMED, Italy). Prior to the walking trials, we recorded a 5-minute standing baseline and used the average of the final 2 minutes as each subject’s baseline metabolic cost. Whole-body kinematics were recorded using a motion capture system (T20, Vicon, UK) at 100 Hz. A 40-marker set was placed on lower-limb segments following a modified Helen Hayes Hospital marker configuration [56].

### 4.2. Predictive simulations of motor performance in novel environments

#### Trajectory optimization model

A standard model-based approach for predicting locomotor performance in novel conditions is trajectory optimization. In this framework, a musculoskeletal model generates movement trajectories (e.g., joint kinematics over a gait cycle) that minimize an experimenter-defined objective function under specified conditions, such as exoskeleton assistance [57], prosthesis use [11], or split-belt walking [12]. This approach yields a steady-state solution that optimizes the chosen cost function, most commonly metabolic energy [58]. We used the 2D musculoskeletal model developed by Price *et al*. [12], which was specifically designed for split-belt walking, as our trajectory optimization benchmark. For each belt speed difference, the model simulated steady-state walking with the corresponding fast and slow belt speeds, generating predictions of metabolic cost (W/kg) and gait symmetry. Because the trajectory optimization framework does not explicitly model adaptation timescales, we focused on the predicted steady-state performance. All simulations were performed in OpenSim 4.3 [59] using OpenSim Moco for trajectory optimization [60]. Each gait cycle was discretized into 100 evenly spaced collocation points and solved using CasADi [61] and IPOPT [62]. Metabolic cost was computed using the muscle activations from the optimized solution and the metabolic model described by Umberger *et al*. [63]. We then used OpenSim to obtain whole-body kinematics, including joint and limb kinematics (e.g., heel position). From these kinematics, we derived model-based step lengths and step length asymmetry using the same definitions as in the experimental analysis.

#### Adaptation-aware model of locomotor performance

We next used a recently proposed adaptation-aware model of locomotor learning (*LocAd*) that qualitatively reproduces a wide range of adaptation phenomena [27]. This model incorporates a two-level hierarchy: (1) a low-level, step-to-step feedback controller responsible for maintaining gait stability, and (2) a high-level policy-gradient reinforcement learner that updates control parameters to improve performance over time. These two layers operate on distinct timescales but interact continuously during adaptation (Figure 2). The original model demonstrated that interactions between energy, stability, and symmetry can shape adaptation trajectories, but did not explain how learning hyperparameters such as the learning rate and symmetry weight might be selected in different environmental settings, influencing adaptive and time-varying performance. Here, we used the adaptation-aware model to study how choices of learning parameters influence multi-objective performance across split-belt settings. To replicate the experimental conditions, we normalized the belt speeds in the *LocAd* model using a dimensionless speed (Froude-like) scaling: each belt’s speed was divided by 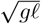, where *g* is gravitational acceleration and *ℓ* is the model’s leg length. This yielded dimensionless slow and fast belt speeds corresponding to each experimental speed difference. To avoid abrupt transitions that frequently produced model falls at large speed differences, we included a 20-second transition period between the tied-belt and split-belt conditions. After each simulation, we computed predicted energy cost based on the mechanical work associated with the step-to-step transition and leg swing [64, 65]. This produced a time-varying energy cost at each stride for each parameter combination and environmental setting.

### 4.3 Inverse adaptation to infer learning parameters and quantify fall risk

#### Aligning simulated and individual-specific time-varying performance

To identify the learning parameters that best explained individual time-varying energetic performance across environmental settings, we implemented an inverse adaptation procedure. For each subject and split-belt condition, we first computed a smoothed experimental energy trajectory by applying a 3-minute moving average to the metabolic cost time series. We then fit this smoothed trajectory with a first-order exponential function and normalized the resulting trend as the percentage change in energy relative to the value at the onset of split-belt walking. We then simulated the adaptation-aware model across a grid of learning rate and symmetry weight values to generate time-varying model energy trajectories. Because the model operates on a stride-by-stride basis rather than continuous time, we applied a 100-stride moving average to the simulated energy to extract the overall trend. We fit this model trend with the same first-order exponential form and normalized it using the same percentage-change convention. For each combination of learning rate and symmetry weight, we ran 50 simulations to account for stochastic variability arising from sensorimotor and exploratory noise in the model. For each subject and condition, we then computed the root mean square error (RMSE) between the experimental and model-derived energy trends. The parameter combination that minimized RMSE was taken as the best-fit learning rate and symmetry weight for that individual and environmental setting. We recorded the best-fit parameters and corresponding RMSE values for all subjects (Figure 3).

#### Effect of model parameters on fall risk across environmental settings

We next used the adaptation-aware model to characterize how learning parameters influence fall risk across environmental settings. Because the model includes stochastic perturbations, repeated simulations with identical parameter values can yield different outcomes (fall or no fall). For each environmental setting and fixed learning rate (or learning rate-symmetry weight pair, as specified below), we therefore performed 50 simulations and quantified fall risk as the fraction of simulations that resulted in a fall. To characterize an additional, more traditional measure of instability, we also computed the number of strides before a fall occurred. During each simulation, we monitored the vertical position of the center of mass; a fall was detected when this height dropped below a predefined threshold (approximating ground level). We then recorded the stride at which the fall occurred. Together, these metrics allowed us to evaluate both probabilistic fall risk and time-to-fall across environmental settings and parameter choices.

#### Visualizing how fall risk explains the shift in learning parameters across environmental settings

To test the hypothesis that the individually fitted learning parameters reflect a fall risk-averse strategy, we examined their relationship to a simulated fall risk landscape. For each belt speed difference, we performed a 2D sweep over learning rate and symmetry weight values and ran 50 simulations per parameter pair. The resulting fall probabilities were used to construct a 2D fall risk surface (heat map) for that environmental setting. We then projected each individual’s best-fit parameter pair (learning rate and symmetry weight) onto the corresponding fall risk landscape. To summarize population-level trends, we computed, for each condition, the 25th and 75th percentiles of the learning rate and symmetry weight distributions and used these to construct an ellipse centered at the median values. Plotting these ellipses atop the fall risk contours for each speed difference allowed us to visualize how parameter distributions shifted across environmental settings and how they related to regions of low versus high fall risk.

### 4.4 Data processing and analysis

#### Metabolic cost

Metabolic cost was calculated using the modified Brockway equation, which incorporates oxygen consumption and carbon dioxide production [66]. The resulting values were normalized by body mass to yield metabolic cost in W/kg. Because breath-by-breath measurements are noisy, we estimated the metabolic cost at each time point by averaging over the preceding 3 minutes of data. Net metabolic cost was obtained by subtracting the standing baseline from the walking metabolic cost.

#### Step length asymmetry

For both experimental and model-based analyses, we identified gait events (heel strikes) for each leg to compute step length and step length asymmetry (SLA). Heel marker positions were tracked in the fore-aft direction, and heel strike events were detected using a local peak detection method (the maximum forward position, corresponding to heel contact). Step length for each leg was defined as the fore-aft distance between the two heels at heel strike. SLA for each gait cycle was then computed as:

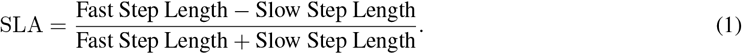

Here, *Fast Step Length* denotes the step length on the fast belt and *Slow Step Length* denotes the step length on the slow belt. A perfectly symmetric gait yields SLA = 0. Negative SLA values indicate longer steps on the slow belt, whereas positive values indicate longer steps on the fast belt.

#### Energy change across environmental settings and timescales

For each subject, we computed the mean metabolic cost over the final 3 minutes of both the tied-belt and split-belt conditions. The effect of the split-belt setting on energy expenditure was quantified as the difference between the split-belt and tied-belt metabolic costs, expressed as a percentage of the tied-belt value to account for inter-individual differences in absolute cost. We then compared these experimental energy changes with predictions from the adaptation-aware model and the trajectory optimization model. For the adaptation-aware model, we computed energy over the final 100 strides of each trial to approximate steady-state behavior. To investigate how adaptation timescales influenced energy changes, we evaluated metabolic cost at 9-minute intervals throughout the 45-minute split-belt exposure. We compared adjacent intervals to assess significant changes and how these varied across belt speed differences (Figure 1E). Because each subject experienced only one split-belt condition, we used a two-way between-subjects ANOVA (factors: environmental setting and time) with a significance level of *α* = 0.05. Significant main or interaction effects were followed by Bonferroni-corrected post hoc pairwise comparisons.

#### SLA change across environmental settings and timescales

We performed a parallel analysis for SLA to characterize how locomotor symmetry changed across environmental settings and timescales. Prior work suggests that changes in body kinematics, including SLA, can precede changes in energy during locomotor adaptation [32]. In split-belt walking, individuals typically exhibit negative SLA early in adaptation, which gradually shifts toward more positive values as they increasingly exploit treadmill work. For each subject, we computed SLA over time and analyzed the change relative to the start of the split-belt condition, using a similar temporal windowing approach as for energy (Figure 1D).

Because SLA is inherently a normalized ratio, we did not further normalize SLA values to baseline. Instead, we calculated the difference in SLA (mean of the last 100 strides) between the split-belt and tied-belt conditions [32, 67]. We then examined how these SLA changes varied across belt speed differences and tested the hypothesis that SLA would increase with larger speed differences, consistent with theoretical predictions that greater mechanical asymmetry should encourage greater reliance on treadmill work.

